# Identifying Alzheimer’s disease-related pathways based on whole-genome sequencing data

**DOI:** 10.1101/2024.07.03.601830

**Authors:** Yongheng Wang, Taihang Liu, Yijie He, Yaqin Tang, Pengcheng Tan, Lin Huang, Dongyu Huang, Tong Wen, Lizhen Shao, Jia Wang, Yingxiong Wang, Zhijie Han

## Abstract

Alzheimer’s disease (AD) is a highly inheritable neurodegenerative disorder for which pathway-specific genetic profiling provides insights into its key biological mechanisms and potential treatment targets. Traditional disease-pathway analyses for AD have certain limitations, such as environmental interference and arbitrary sample division. We present a comprehensive framework that starts with genome data, avoiding these drawbacks and offering intrinsic pathway-specific genetic profiling for AD. Whole genome sequencing data from 173 individuals were used to quantify transcriptomes in 14 brain regions, estimate individual-level pathway variant scores, and analyze AD risk for each patient. These results were combined to identify AD-related pathways and quantify their interactions. The predicted expression levels were consistent with previous findings, and the estimated AD risk showed a significant correlation with Braak/Thal scores. A total of 3,798 pathways were identified as potentially associated with AD, with about 19.7% previously reported. Key pathways, including NF-κB signaling and GSK3β activation, were linked to AD pathogenesis. The interactions among pathways highlighted shared gene functions in AD. In summary, we provided an effective framework for disease-pathway analysis, revealing the interdependence or compensatory effects of pathways in AD.

## Introduction

Alzheimer’s disease (AD) is a progressive neurodegenerative disorder with a high heritability estimated between 60% and 80%^1^. Despite identifying numerous genetic variants associated with AD risk and pathology^2–5^, the precise pathogenesis of AD remains unclear, and effective clinical treatments are still lacking. Pathway-specific genetic profiling provides a means to understand the key biological mechanisms involved in diseases and enables the identification of causal genes and potential therapeutic targets^6^. Therefore, studying the genetic characteristics of AD-related pathways is essential. Typically, the protocol for inferring disease-related pathways relies on detecting differential characteristics between disease and control groups. For example, a case-control-based genotype-phenotype association analysis highlighted the involvement of amyloid/tau pathways in AD^7^. Furthermore, analyzing transcriptomic^8, 9^ and proteomic^10, 11^ differences between AD patients and control individuals has discovered several new AD-related pathways.

However, the case-control design has limitations: (1) These analyses require artificially dividing samples into disease and control groups. For diseases like AD, which exhibit a progressive state of risk and severity, it is challenging to establish a clear standard for distinguishing whether a person is diseased or not. For example, early AD diagnosis relies on assessing AD neuropathological changes, including neurofibrillary tangle distribution (Braak NFT stage), amyloid-beta plaques (Thal stage), and the neuronal plaques density in the neocortex (CERAD score). In the preclinical (asymptomatic) phase of the AD continuum, misdiagnosis and missed diagnosis may occur^12^. Additionally, the observed phenotype of AD patients (e.g., behavioral rating scale) and their pathological diagnosis (e.g., senile plaque and neurofibrillary tangle pathology) can sometimes be contradictory^13^. (2) Analysis usually starts with quantifying expression levels, which can be influenced by environmental factors, leading to potentially false associations with disease pathogenesis. This includes incorrect findings of blood and cerebrospinal fluid-based biomarkers used for early AD diagnosis^14^. (3) Even considering environmental factors, determining disease-related genes and pathways is often based on a one-size-fits-all threshold, which may result in losing a significant amount of potentially relevant gene information.

Genomic information provides valuable insights into the original genetic characteristics and the molecular basis of pathogenesis. Disease-related single nucleotide polymorphisms (SNPs), genes, and individual disease risks can be identified from genomic data using methods like genome-wide association studies (GWAS), fine-mapping, linkage disequilibrium score regression (LDSC), and polygenic risk score (PRS)^15, 16^. Compared to single SNP analyses, gene and gene set analyses are considered more powerful^17^. Gene set analysis, in particular, offers valuable insights into the role of specific biological pathways or cellular functions in the genetic basis of phenotypes^18^. While pathway-based PRS tools like PRSet^19^, MAGMA^18^, and stratified LDSC^17^ exist, they typically only assign SNPs to genes, then genes to gene sets, and construct test statistics. These tools link variants to pathways based on their location, without considering the correlation between SNPs and gene expression levels, characterized by expression quantitative trait loci (eQTL).

To address these limitations, this study developed a new framework to investigate the genetic characteristics of AD pathways directly from the genome. Our framework identified pathways related to AD in different brain regions based on their genetic characteristics. The advantages of our analysis are noteworthy: it requires only genotyping data (obtaining brain tissue is challenging, but blood samples are easily accessible), and no sample grouping information is necessary. Additionally, this framework can analyze continuous phenotype variables. By starting our analysis directly from the genome, we achieve results closer to the genetic characteristics. We believe this framework holds promise for providing quantitative insights into the combined effects between pathways in complex diseases in the future.

## Results

### Pipeline overview

To investigate the genetic characteristics of AD, we used whole genome sequencing (WGS) data from 173 individuals (including 92 AD cases and 81 controls according to the original annotations) obtained from the MayoRNAseq (https://doi.org/10.7303/syn10901601). Since most associations between genes/pathways and diseases are tissue-specific^20^, we first used a method called genetically regulated component of expression (GReX) to produce genome-wide expression data from 14 brain regions of GTEx (v8)^21^. This genotype data-based approach eliminates potential confounding factors that affect gene expression levels, such as environmental influences, allowing us to focus on the genetic regulation^22^. We then used the quality-controlled genotype data to estimate individual AD risk scores by PRS, instead of categorizing individuals based on original annotations. Subsequently, we performed a single sample gene set variation analysis (ssGSVA) to analyze individual-level pathway variant scores based on quantified gene expression^23^. We combined these scores with PRS to identify AD-related pathways through a distance rank correlation analysis^24^. Finally, we conducted a structural equation modeling (SEM) analysis to quantify interactions among the AD-related pathways, identifying possible genetic etiology and pathway mechanisms of AD (**Figure 1**).

**Figure 1.**
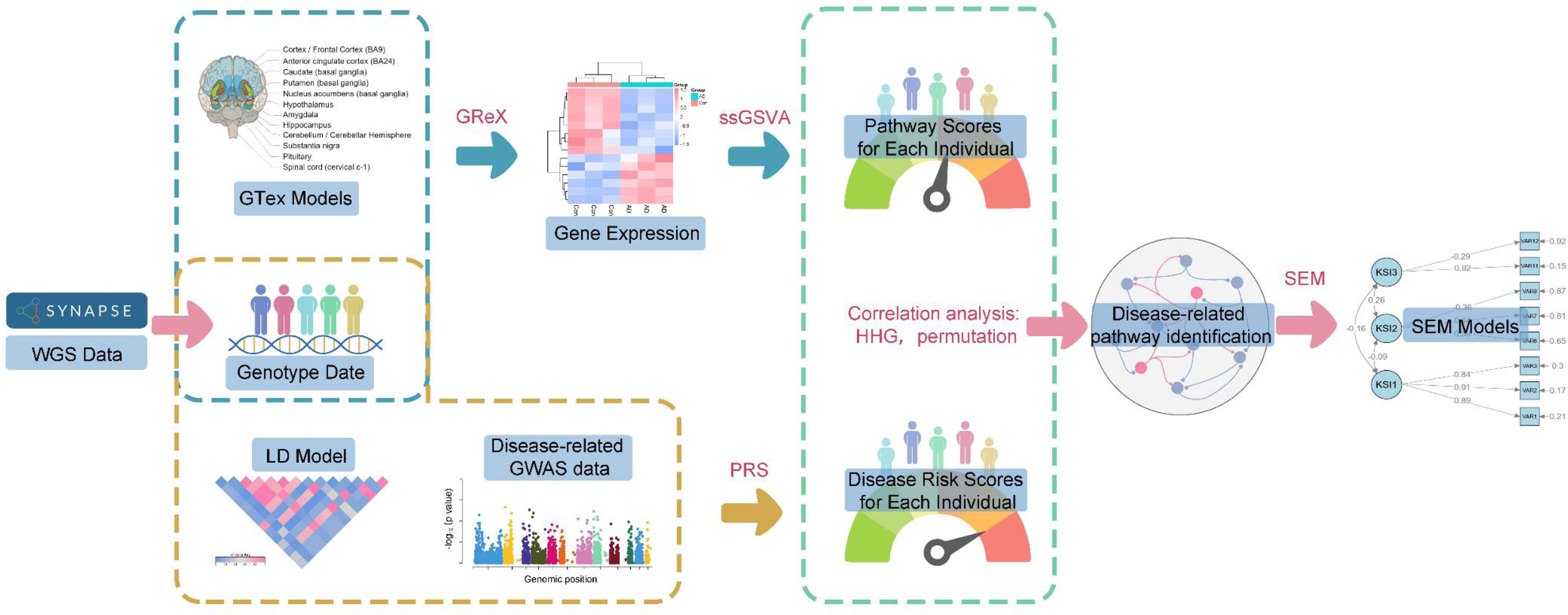
Pipeline schematic of investigating the original genetic characteristics of AD pathways. In this pipeline, only genotype data is required, which can be obtained through various methods such as WGS, whole exome sequencing (WES), or SNP array analysis. Pathway scores and disease risk scores for each individual were calculated based on their genotype data. Subsequently, analyses were conducted to explore AD-related pathways and the interaction models between these pathways. This genotype data-driven approach eliminates potential confounding factors that influence gene expression levels, allowing for a focused investigation into genetic regulation. Furthermore, this approach can also be adapted for analyzing genetic regulatory pathways in other diseases.

### Transcriptome prediction and gene sets variation analysis based on genotype data

To avoid the interference of environmental factors, we predicted the gene expression from the genomic data by GReX method^22^. From the 349 samples in the MayoRNAseq WGS study, we selected 92 AD samples and 81 control samples, all of white race. Based on the genomic features, we predicted transcriptome expression in 14 brain regions from GTEx (v8), including: amygdala, anterior cingulate cortex BA24, caudate basal ganglia, cerebellar hemisphere, cerebellum, cortex, frontal cortex BA9, hippocampus, hypothalamus, nucleus accumbens basal ganglia, putamen basal ganglia, spinal cord cervical c-1, substantia nigra, and pituitary. As shown in **Figure 2A**, the number of predicted genes varied, ranging from 5,013 in substantia nigra to 9,186 in cerebellum. The predicted gene expression levels ranged from −3 to 3. Furthermore, pairwise correlation analysis of brain regions revealed that gene expression correlations between the cerebellum, cerebellar hemisphere, and pituitary, and other brain regions were all below 0.85 (**Figure 2B**). In contrast, correlations between other brain regions were relatively high. The correlation between similar brain regions exceeded 0.93, including the caudate basal ganglia, nucleus accumbens basal ganglia, and putamen basal ganglia; the cortex and frontal cortex BA9; and the cerebellar hemisphere and cerebellum. These results align with previous findings that gene expression regulation shows specificity in the cerebellum and similarity across various regions of the cerebral cortex^25^.

**Figure 2.**
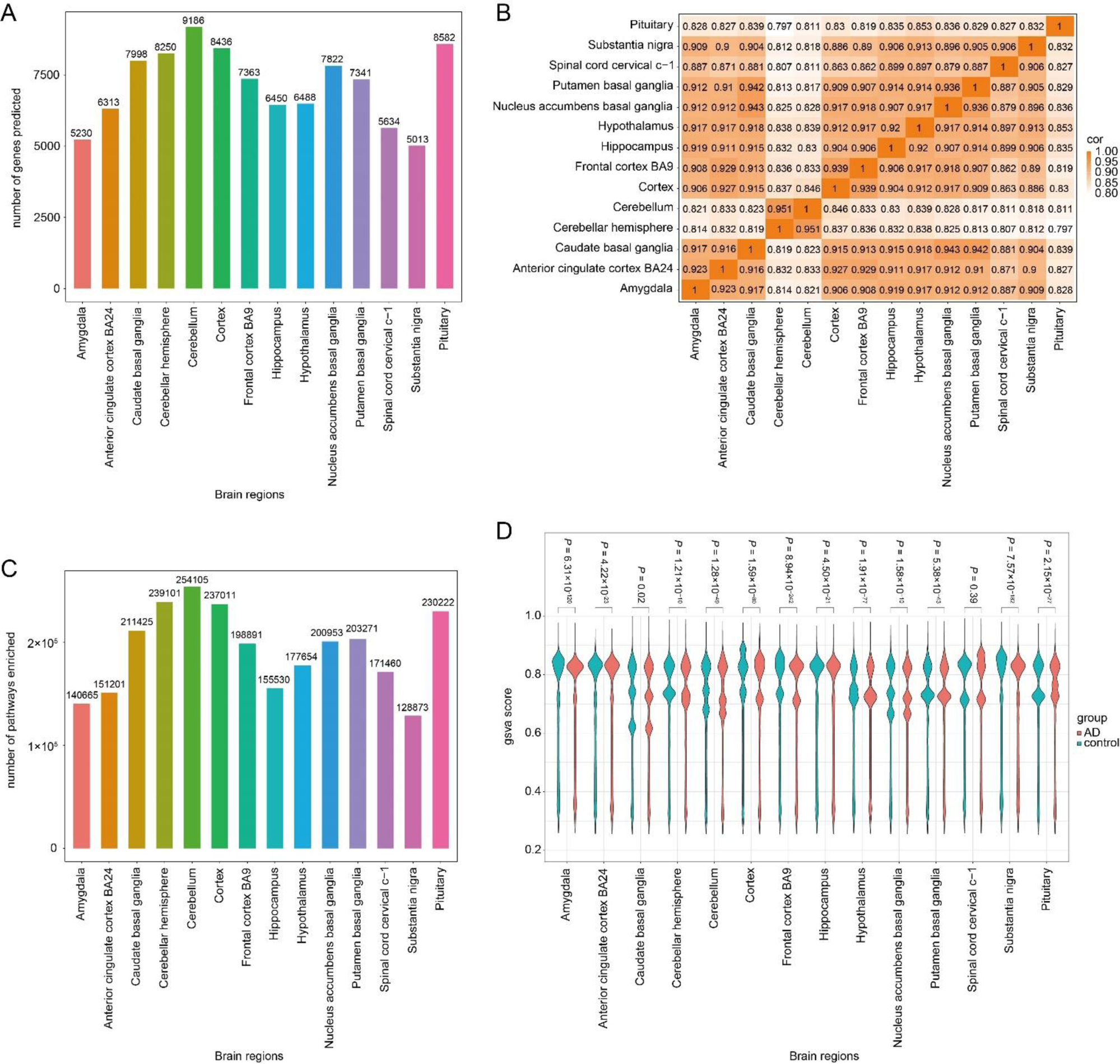
Performance of GReX and ssGSVA. (**A**) The number of genes predicted by GReX method for the 14 brain regions. (**B**) The pairwise correlations computed using the Pearson method among 14 brain regions based on the predicted expression levels. (**C**) The number of pathways with ssGSVA score enriched by ssGSVA method for the 14 brain regions. (**D**) The distribution of ssGSVA scores in AD and control samples.

To identify AD-related pathways, we used the non-parametric and unsupervised ssGSVA method^23^. For the ssGSVA analysis, we integrated gene sets of 1,649,335 pathways from multiple resources, including GSEA^26^, DAVID^27^, Reactome^28^, NetPath^29^, PANTHER^30^, WikiPathways^31^, and PathBank database^32^. We then calculated ssGSVA scores for each individual across 14 brain regions. As shown in **Figure 2C**, the number of pathways with ssGSVA score ranged from 128,873 in the substantia nigra to 254,105 in the cerebellum, which consistent with the trend observed in the number of predicted genes. Additionally, we calculated the average ssGSVA score for each pathway among the originally annotated AD cases and controls, respectively, and compared their distribution in the 14 brain regions using a two-tailed Wilcoxon test (the threshold of *P* < 0.05). We observed significant differences in most brain regions, except the spinal cord cervical c-1 (*P* = 0.39) (**Figure 2D**).

### Identification of AD risk associated pathways

To estimate the genetic liability to AD risk in individuals, we performed PRS analysis. Following PRS guidelines, we retained 173 samples (92 AD cases and 81 controls according to the original annotations) and 10,070,779 SNPs after applying filters and quality control (QC). By combining with the QCed AD GWAS summary data, we calculated the AD risk score for these 173 individuals. We assessed the similarity between the AD risk scores and the original annotations by performing linear Pearson’s analysis. Despite some differences, the AD risk score showed a significant correlation with individual group (*R* = 0.48, *P* = 3.54×10^−11^), Thal stage (*R* = 0.54, *P* = 3.39×10^−11^), and Braak NFT stage (*R* = 0.56, *P* = 5.25×10^−12^) (**Figure 3A**). We also compared the distribution of the average ssGSVA scores between individuals with high (more than zero) and low (less than zero) AD risk scores in the 14 brain regions using the two-tailed Wilcoxon test, respectively. Significant differences were found in all 14 brain regions (*P* < 0.05), and these differences were more pronounced overall compared to those observed in the previous step (grouping by original annotations) (**Figure 3B**).

**Figure 3.**
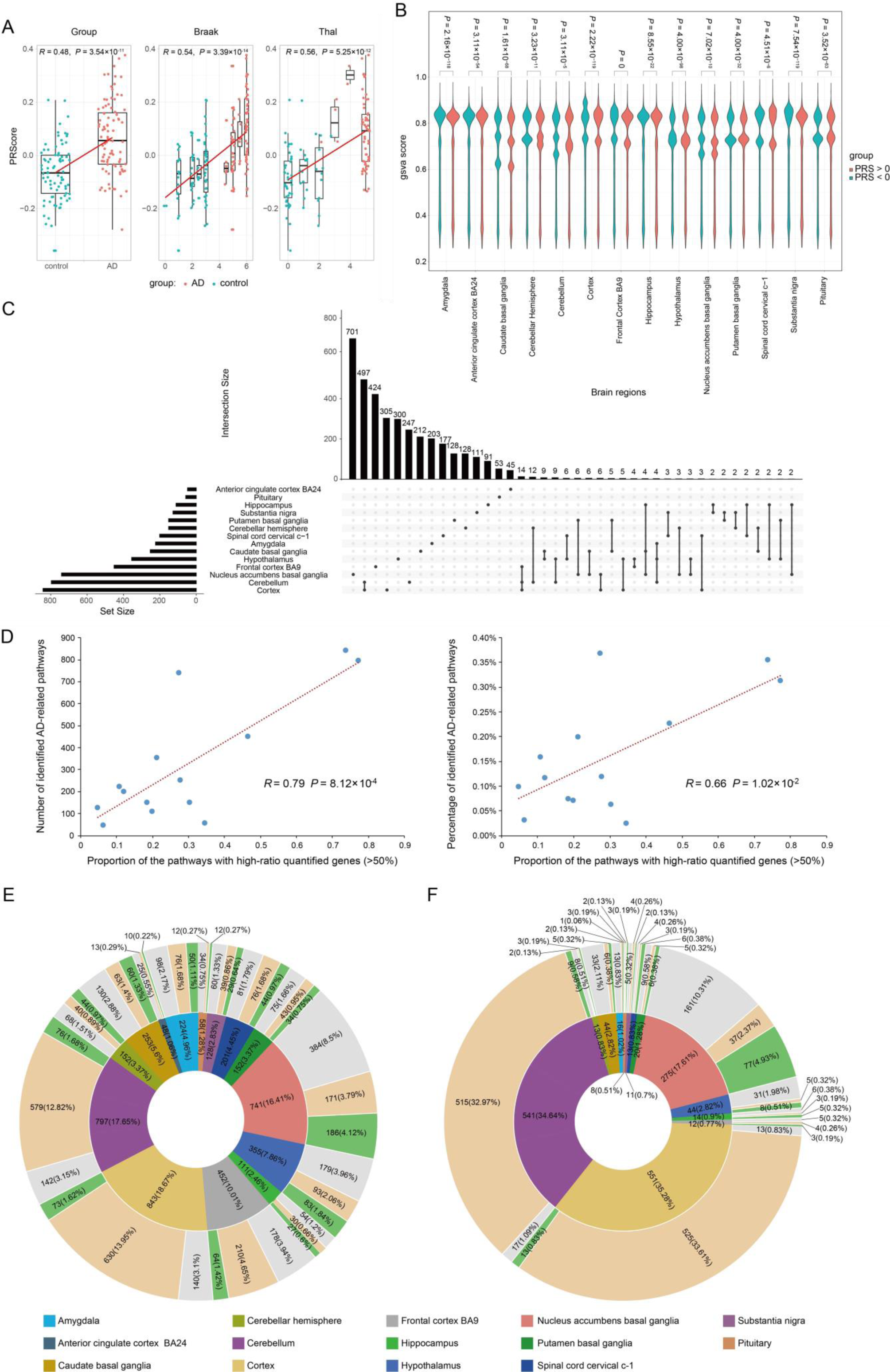
Performance of PRS and identification of AD-related pathways. (**A**) Spearman correlation analysis depicting the association between AD risk score (measured by PRS analysis) and AD/control group, Braak stage, and Thal stage. (**B**) The distribution of ssGSVA scores in high (more than zero) and low (less than zero) AD risk samples. (**C**) Visualization of intersecting sets of AD-related pathways identified by HHG analysis and permutation procedure. (**D**) The relationship between the proportion of pathways with high-ratio quantified genes and the efficiency of AD-related pathway identification. (**E**) The proportion of direct (green), indirect (burlywood) and unrelated (grey) pathways within each brain region among all AD risk-related pathways and (**F**) among all pathways in SEM results.

Further, we identified a total of 3,798 pathways significantly associated with the AD risk score. Since most ssGSVA scores do not follow a normal distribution (Shapiro-Wilk test, *P* < 0.05), and the correlation pattern between ssGSVA scores and AD risk scores was unknown, we employed the Heller-Heller-Gorfine (HHG) method to assess the correlation^33, 34^. Before conducting the correlation analysis, we removed outliers from the scores. **Table 1** demonstrates that the number of pathways significantly related to AD risk in each brain region, identified through a permutation procedure, ranged from 48 (anterior cingulate cortex BA24) to 843 (cortex). Moreover, pathways were identified in at least two brain regions accounting for about 16% of the total (**Figure 3C**). It is worth noting that the significant correlation pathways identified by HHG (*P* < 0.001) matched those identified by the permutation procedure. In addition, we define a pathway in which more than 50% of genes are quantified by GReX and used for ssGSVA scoring as a pathway with high-ratio quantified genes. We observed that the more pathway-related genes quantified in a tissue, the more AD-related pathways were identified within it. **Figure 3D** shows a significant positive correlation between the proportion of pathways with high-ratio quantified genes and the percentage of identified AD-related pathways (*R* = 0.79, *P* = 8.12 ×10^−4^). This suggests that more comprehensive eQTL data can enhance the efficiency of our framework.

**Table 1.** Overview of the genes and pathways predicted in 14 brain regions.

| Brain region | Pathway |  |  |  | Gene |  |  |
| --- | --- | --- | --- | --- | --- | --- | --- |
|  | ssGSVA | AD risk-related | literature-reported AD-related | Percentage<sup>a</sup> | predicted | In literature-reported AD pathways | Percentage<sup>b</sup> |
| Amygdala | 140665 | 224 | 50 | 22.32% | 5230 | 1038 | 19.85% |
| Anterior cingulate cortex BA24 | 151201 | 48 | 10 | 20.83% | 6313 | 160 | 2.53% |
| Caudate basal ganglia | 211425 | 253 | 59 | 23.32% | 7998 | 1015 | 12.69% |
| Cerebellar hemisphere | 239101 | 152 | 44 | 28.95% | 8250 | 1007 | 12.21% |
| Cerebellum | 254105 | 797 | 76 | 9.54% | 9186 | 906 | 9.86% |
| Cortex | 237011 | 843 | 72 | 8.54% | 8436 | 4078 | 48.34% |
| Frontal cortex BA9 | 198891 | 452 | 64 | 14.16% | 7363 | 786 | 10.67% |
| Hippocampus | 155530 | 111 | 27 | 24.32% | 6450 | 431 | 6.68% |
| Hypothalamus | 177654 | 355 | 83 | 23.38% | 6488 | 2494 | 38.44% |
| Nucleus accumbens basal ganglia | 200953 | 741 | 187 | 25.24% | 7822 | 1567 | 20.03% |
| Putamen basal ganglia | 203271 | 152 | 34 | 22.37% | 7341 | 651 | 8.87% |
| Spinal cord cervical c-1 | 171460 | 201 | 43 | 21.39% | 5634 | 398 | 7.06% |
| Substantia nigra | 128873 | 128 | 29 | 22.66% | 5013 | 254 | 5.07% |
| Pituitary | 230222 | 58 | 12 | 20.69% | 8582 | 109 | 1.27% |

Percentage<sup>a</sup>: number of literature-reported AD-related pathways/number of AD risk-related pathways. Percentage<sup>b</sup>: number of genes in literature-reported AD-related pathways/number of predicted genes.

### Literature reported relation between AD risk-related pathways and AD

We investigated the literature-reported correlation between the 3,798 AD risk-related pathways identified across 14 brain regions and AD. These correlations were classified as direct, indirect (associated with features linked to AD), or unrelated. **Table 1** provides the number of direct AD risk-related pathways, pathways directly associated with AD with supporting literature, and the percentage of direct correlations within each brain region. Detailed descriptions of the reported relevance between the enriched pathways and AD are in **Supplementary Table S1**, along with corresponding references. Additionally, **Figure 3E** illustrates the proportion of direct, indirect, and unrelated pathways within each brain region among all AD risk-related pathways. The pathways directly correlated with AD mainly include those related to the immune system, regulation of Aβ, APP, tau, and other key genes/proteins, autophagy, mitochondrial/endoplasmic reticulum (ER)-mitochondria interactions, mTOR, NFkappaB, estrogens/androgens, helicase/DNA repair, HLA, metal ion metabolism, miRNAs, ROS, p38 MAPK, and VEGF.

The number of predicted genes, the genes in these pathways directly associated with AD, and the percentages are also listed in **Table 1**. Two brain regions have significantly higher proportions of gene counts: cortex (48.34%) and hypothalamus (38.44%). This is due to the presence of two AD-related pathways in each of these brain regions that contain an unusually high number of genes. Specifically, the total number of genes included in the pathways used for ssGSVA is 44,659, with each of these four pathways containing more than 4,000 genes. In the cortex, two AD-related pathways (GOBP ∼ GO:0008152 ∼ metabolic process and GOBP ∼ GO:0071704 ∼ organic substance metabolic process) originally contain a high number of genes (12,193 and 11,793, respectively), and 3,820 and 3,664 of them were used in ssGSVA, respectively. Similarly, in the hypothalamus, two AD-related pathways (UCSC_TFBS ∼ TBP and UCSC_TFBS ∼ CEBPA) also originally contain a high number of genes (5,762 and 4,661, respectively), and 1,235 and 977 of them were used in ssGSVA, respectively. The rest of the pathways in these two brain regions contain several to several tens of genes (totaling 1,058 in the cortex and 1,220 in the hypothalamus), which is similar to other brain regions. Additionally, the nucleus accumbens basal ganglia has the highest proportion of AD-related genes (20.03%).

### Quantitatively investigate the interaction relationships among pathways

Given that pathways do not work independently^35, 36^, we utilized SEM analysis to quantitatively assess their interactions and contributions to the phenotype. This process involved factor analysis, model building, and optimization using AD risk-related pathways with unreplicated ssGSVA scores. As shown in **Table 2**, the Bartlett test for all 14 brain regions was significant after removing variables with inadequate loadings and those with only one variable in a factor. Additionally, KMO values exceeded 0.5 for most brain regions (except for the anterior cingulate cortex BA24, frontal cortex BA9, and pituitary), indicating the appropriateness of factor analysis. After model optimization, the standard error of pathway loading in all brain regions within their respective factors was significant (generally t value > 2), except for the anterior cingulate cortex BA24. The modification indices for pathways in other factors remained relatively low, indicating a generally well-fitting model. Except for the anterior cingulate cortex BA24, the key results of the SEM models for the other 13 brain regions (including pathway attribution factors, loadings, and inter-factor correlations, as well as the corresponding pathways and their associations with AD) can be found in **Supplementary Table S2** to **S14**. **Table 2** lists the number and proportion of literature-reported AD-related pathways in each brain region model. Consistent with the proportion of gene counts (**Table 1**), the model for the nucleus accumbens basal ganglia also contains the largest number of AD-related pathways. Additionally, **Figure 3F** shows the proportion of direct, indirect, and unrelated pathways within each brain region among all pathways in the SEM results. **Figure 4** provides a detailed overview of the SEM results for the nucleus accumbens basal ganglia, grouping pathways with similar functions into the same factor, regardless of their originating databases. For instance, four pathways, including “anti-inflammatory actions mediated by gliotoxin include HO-1 induction and the subsequent blockade of NF-kappaB-dependent signaling pathways in vitro and in vivo (GENERIF_SUMMARY∼16804400)”, “GA could inhibit NF-kappaB and MAPK/HO-1 signalling pathways (GENERIF_SUMMARY∼24300974)”, “induction of HO-1 by CO-RM2 exerted anti-inflammatory and antioxidant effects which are required in concert to prevent the activation of NF-kappaB leading to induction of various inflammatory genes implicated in the pathogenesis of RA (GENERIF_SUMMARY∼24616552)”, and “Induction of Prx1 by hypoxia regulates heme oxygenase-1 via NF-kappaB in oral cancer (GENERIF_SUMMARY∼25162226)” belong to factor KSI8 within the NFkappaB category. These pathways share the genes HMOX1 (ENSG00000100292) and NFKB1 (ENSG00000109320), which interact and affect AD risk^37^. The inter-factor correlations in this brain region can be found in **Supplementary Table S10**.

**Figure 4.**
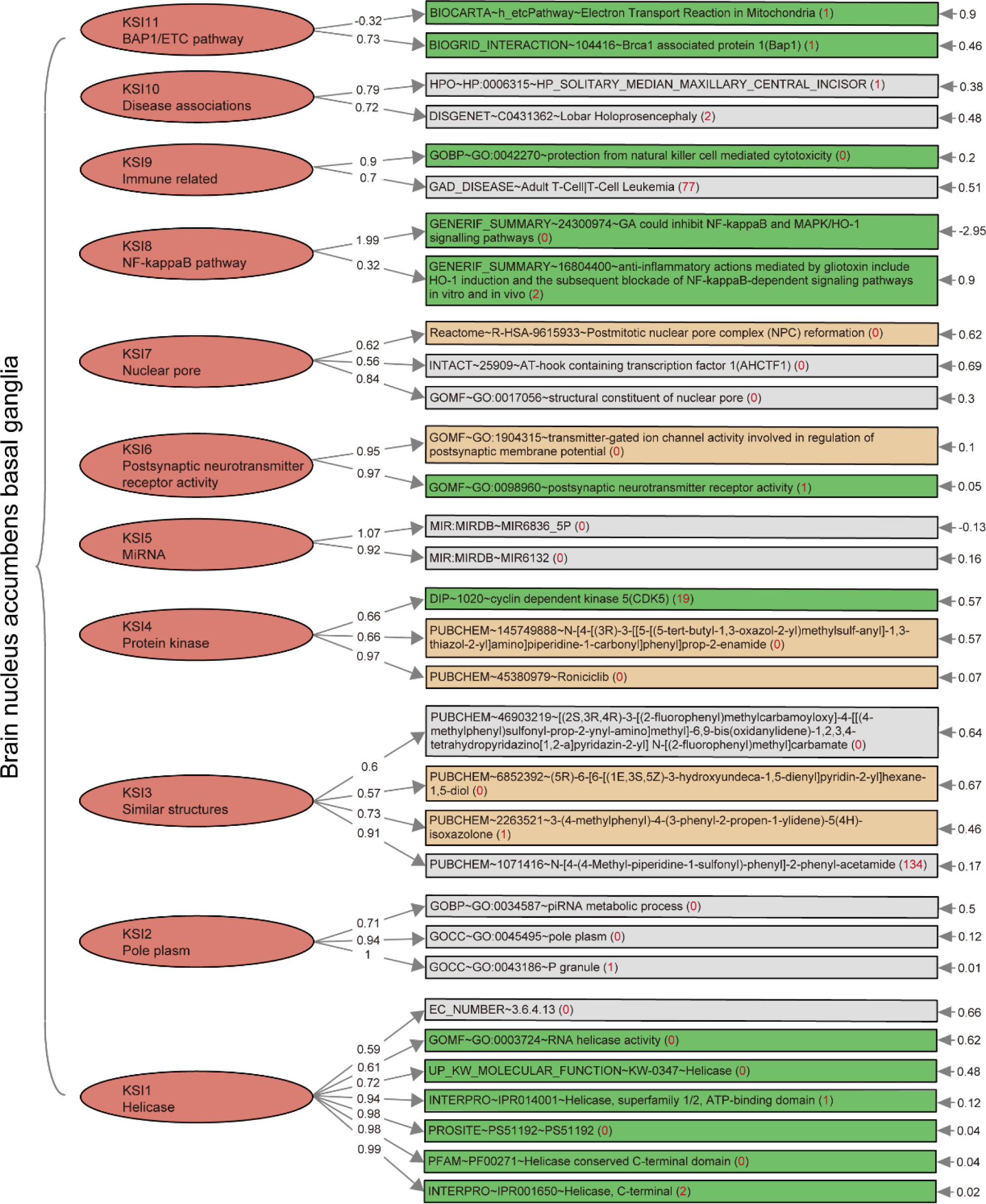
The SEM result of nucleus accumbens basal ganglia. The red numbers in parentheses represent number of pathways with the same ssGSVA score as this pathway in this model. The inter-factor correlations in this brain region can be found in **Supplementary Table S10**.

**Table 2.** SEM analysis results of 14 brain regions.

| Brain region | KMO | Bartlett | Number of pathways in module | Number of factors | Number of literature-reported AD-related pathways | Percentage |
| --- | --- | --- | --- | --- | --- | --- |
| Brain amygdala | 0.516 | 0.000 | 16 | 5 | 1 | 6.25% |
| Brain anterior cingulate cortex BA24 | 0.483 | 0.000 | NA | NA | NA | NA |
| Brain caudate basal ganglia | 0.588 | 0.000 | 44 | 9 | 5 | 11.36% |
| Brain cerebellar hemisphere | 0.615 | 0.000 | 13 | 3 | 3 | 23.08% |
| Brain cerebellum | 0.617 | 0.000 | 541 | 9 | 9 | 1.66% |
| Brain cortex | 0.632 | 0.000 | 551 | 9 | 13 | 2.36% |
| Brain frontal cortex BA9 | 0.493 | 0.000 | 12 | 5 | 3 | 25.00% |
| Brain hippocampus | 0.539 | 0.000 | 14 | 6 | 5 | 35.71% |
| Brain hypothalamus | 0.607 | 0.000 | 44 | 11 | 8 | 18.18% |
| Brain nucleus accumbens basal ganglia | 0.649 | 0.000 | 275 | 11 | 78 | 28.36% |
| Brain putamen basal ganglia | 0.506 | 0.000 | 20 | 8 | 6 | 30.00% |
| Brain spinal cord cervical c-1 | 0.516 | 0.000 | 13 | 6 | 6 | 46.15% |
| Brain substantia nigra | 0.506 | 0.000 | 11 | 4 | 2 | 18.18% |
| Pituitary | 0.499 | 0.000 | 8 | 4 | 3 | 37.50% |

## Discussion

In this study, we predicted transcriptomes of 14 brain regions based on genotype data. We then combined these transcriptomes with GWAS summary data to estimate the PRS for AD risk. Subsequently, we characterized the primary inheritance pathways associated with AD. This framework for identifying disease-related pathways offers three main advantages: (1) By starting from genome data, we avoided environmental factors’ interference, yielding results that more accurately reflect the disease’s intrinsic genetic mechanisms. (2) Our approach relied solely on genotype data, eliminating the need to divide samples into disease or control groups. This enabled us to analyze continuous variables, which provide a more nuanced representation of the real-world disease landscape. (3) Our research covered the entire genome without imposing an arbitrary threshold, preventing the loss of important genomic information.

The number of genes predicted by GReX is relatively high in the cerebellum/cerebellar hemisphere and pituitary, but their correlation with other brain regions is low. This is consistent with the eQTL results, indicating a higher prevalence of genes exhibiting allele-specific expression in these tissues^21, 38^. The trend of pathway numbers enriched by ssGSVA aligned with the predicted gene numbers. The PRS reflected the genetic risk for AD and showed a significant correlation with AD/control grouping, as well as the Braak and Thal stages of the samples. Overall, the higher the Braak and Thal stages, the greater the AD risk. Through correlation analysis, we identified pathways associated with AD risk in each brain region. Of these pathways, 19.7% have been previously reported to be related to AD, while 39.7% may potentially be associated with AD. Both PRS and ssGSVA scores are derived from genomic data, allowing us to elucidate AD-related pathways purely from a genetic perspective. Further, the SEM analysis enabled us to assess the interaction and coefficient among pathways. Pathways assigned to one factor tended to share more genes and exhibit similar functions.

Research on AD encompasses various aspects such as immune regulation^39^, autophagy^40, 41^, mitochondrial function^42^, and more. The AD-related pathways identified across 14 brain regions span multiple domains, highlighting their significance. As illustrated in **Figure 5**, we have summarized a regulatory network possibly involved in the regulation of AD by consolidating several AD-related pathways that we have identified. Specifically, the transcription factor NF-κB signaling can be activated through the TLR4/TRAF6/IKK or PI3K/AKT signaling pathways, promoting the transcription of cytokines such as IL-1, TNF-α, and chemokines like MCP-1^43, 44^. This activation stimulates neuroinflammation in cells such as microglia, contributing to the pathogenesis of AD. Amyloid β can transmit signals by binding to its membrane receptor PirB, which then activates GSK3β via PP2A^45^. GSK3β, acting as a multifunctional factor, promotes the phosphorylation of tau and MACF1, leading to their detachment from microtubules, resulting in abnormal microtubule structures and neuronal damage^45, 46^. Additionally, it promotes the degradation of β-catenin by phosphorylation, inactivating the Wnt signaling pathway and impairing neuronal development and function^47, 48^. HSP90 binds to phosphorylated tau and restores its normal structure by dephosphorylation. Abnormal ubiquitination of the protein c-Cbl is closely associated with AD, as it can ubiquitinate and degrade proteins such as AKT, NF-κB, β-catenin, and HSP90^49^, all of which are involved in the occurrence of AD.

**Figure 5.**
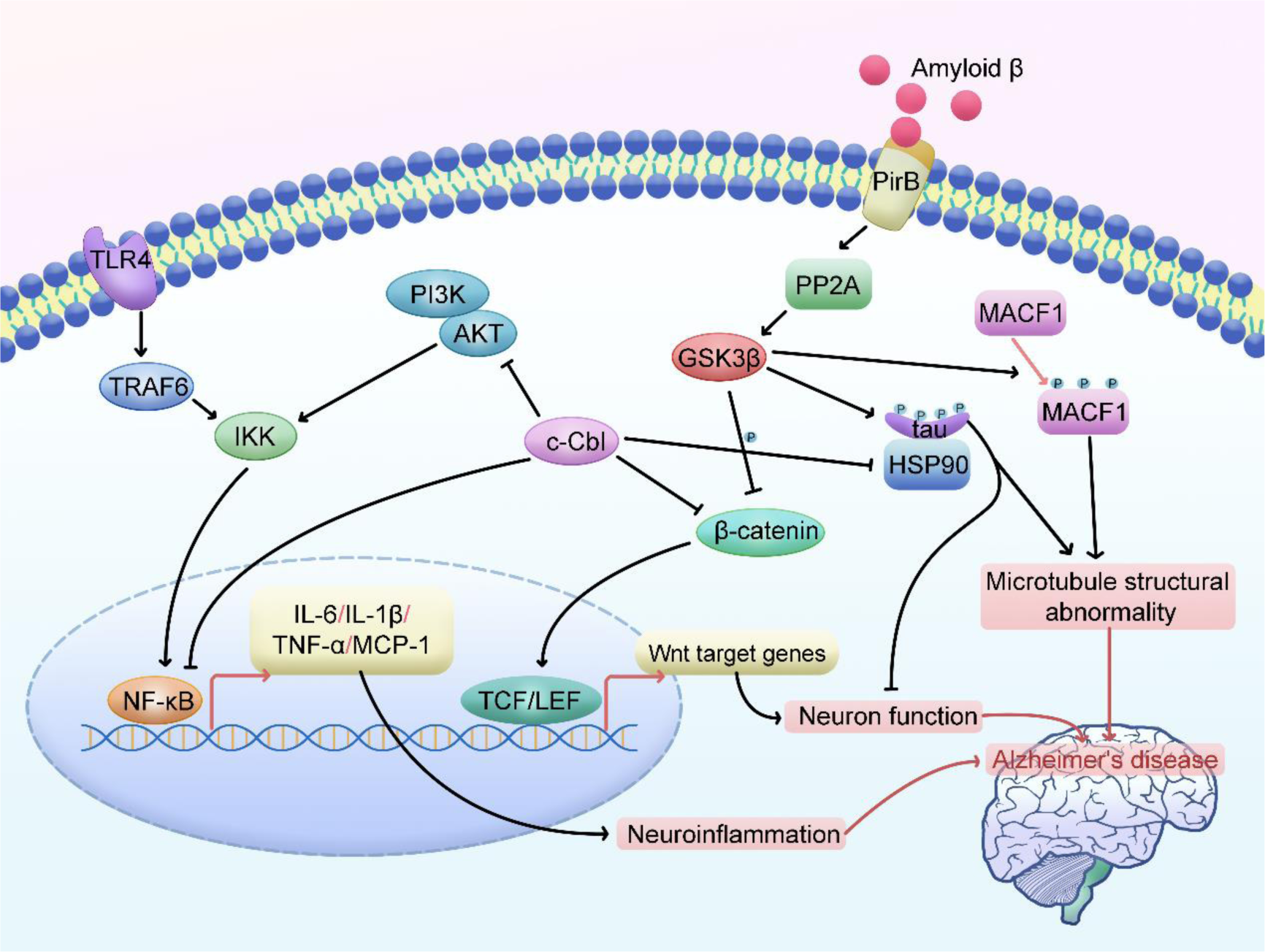
A pathway network summarized by consolidating several AD-related pathways that we have identified.

In conclusion, given the numerous alterations observed in various pathways in AD, targeting a single process will likely prove insufficient to halt disease progression. More importantly, our findings have pinpointed specific categories that could serve as a reference for investigating pathway-dependent AD risk, offering valuable quantitative insights into the interdependence or compensatory effects among pathways. Further research is warranted to explore the precise roles of pathways and their interrelationships with AD.

## Methods

### Whole genomic data

The WGS data from the MayoRNAseq study were utilized in this study. All samples passed data QC as described. Based on the phenotypic information provided, the samples were diagnosed according to the NINCDS-ADRDA criteria and the neuropathologic evaluation^50^. All individuals diagnosed with AD had an empirical diagnosis and were in Braak NFT stage IV or greater. For the control subjects, their Braak NFT stage was III or less, and they exhibited CERAD neuritic and cortical plaque densities of 0 (none) or 1 (sparse). Additionally, the control group did not have any other pathological diagnosis that could potentially confound the AD-related results. We selected samples with a race of white and a diagnosis of AD or control for the following analysis. The genotype profile data of the selected samples were then processed into the format required for gene expression prediction and PRS analysis using plink 1.9^51^.

### Genome-wide gene expression prediction

The gene expression patterns in different brain regions of individuals were assessed based on GReX approach from the MetaXcan project (https://github.com/hakyimlab/MetaXcan)^22^. To achieve this, we combined the genotype data of MayoRNAseq and eQTL data of GTEx (v8) to establish associations between gene expression and genetic variation in 14 brain regions through linear regression modeling. Subsequently, we used the GReX model with its default parameters to calculate the gene expression in these brain regions. Pairwise correlations between brain regions were calculated using the average gene expression for each region.

### Gene-pathway association collection and ssGSVA

We integrated the mapping relationships between pathways and genes from several authoritative databases. Since the pathway data were collected in various formats, we converted them into the list format necessary for conducting ssGSVA. First, we filtered out pathways from various databases that encompassed multiple species, including PathBank (https://pathbank.org/)^32^, DAVID (v.2022q2) (https://david.ncifcrf.gov/)^27^, NetPath (http://netpath.org/)^29^, Reactome (v81) (https://reactome.org/)^28^, WikiPathways (v.20220710) (https://classic.wikipathways.org/index.php/)^31^, GSEA (MSigDB v7.5.1) (https://www.gsea-msigdb.org/gsea/index.jsp)^26^, and PANTHER (http://www.pantherdb.org/)^30^. Only the human pathways were exclusively selected for further analysis. Second, all gene names or IDs were converted to Ensembl gene IDs. Pathway names were organized in the format of type ∼ ID ∼ term (e.g., GOBP ∼ GO:0000002 ∼ mitochondrial genome maintenance). Some pathways were in type ∼ term format if they did not have IDs. Additionally, the same pathway in different databases was updated to the latest version. Finally, the curated data were organized into lists of gene sets for ssGSVA. We integrated the pathway information into the Rdata format and stored them at https://github.com/BFGBgroup/Pathway/blob/main/geneSets.Rdata. The R package ‘GSVA’ was used to calculate the score of each pathway in different brain regions for individuals. Considering that a gene set (pathway) has an ssGSVA score of 1 in all samples when only one gene expression is detected in its expression matrix, we set the parameter min.sz (minimum number of genes) to 2.

### Polygenic risk score (PRS) analysis

The GWAS summary statistics data used in this study were obtained from the research of Jansen *et al.*, including 71,880 AD cases and 383,378 controls^52^. The data were downloaded from a public database (https://ctg.cncr.nl/software/summary_statistics/). For PRS analysis, standard quality control of GWAS summary data and genotype profile data was performed according to the guidelines for PRS analysis^53^. PRScs was then used to calculate the posterior SNP effect size for each chromosome, and the output files from all chromosomes were concatenated^54^. The 1000 Genomes Project phase 3 European subset was employed as the LD reference panel^55^. Finally, the individual-level polygenic score was generated using plink 1.9^51^. Correlations between PRS and phenotypes (AD/control, Braak NFT stage, Thal) were calculated using the Spearman method.

### Correlation analysis between PRS and ssGSVA score

The Shapiro-Wilk test was used to determine if the ssGSVA scores of pathways followed a normal distribution. The R package ‘HHG’ was employed to calculate the correlation between ssGSVA scores and PRS^33, 34^. Outliers were removed using the ‘winsor’ function of the R package ‘psych’ before calculating correlations. To identify pathways significantly associated with the PRS, a permutation procedure was used to correct for the multiple hypothesis testing effect from analyzing numerous pathways. Specifically, permutations were performed by randomizing the PRS and ssGSVA scores. Between 1,000 and 10,000 permutations were conducted for each pathway’s correlation analysis with PRS. The stopping condition was set to ensure that at least 15 permuted *P* values were smaller than the nominal *P* value within this range. The threshold for each pathway was determined as follows: First, the empirical *P* value of each pathway was defined as the ratio of occurrences of extreme events (permuted *P* values smaller than the nominal *P* value) to the total number of permutations. Then, for each brain region, the empirical *P* values were corrected to empirical q-values using Storey’s approach. The empirical *P* value that corresponded to the empirical q-value closest to 0.05 was selected. Finally, the threshold for each pathway was set as the nth permuted *P* value in ascending order, where n represents the ceiling integer of multiplying the selected empirical *P* value for the brain region by the number of permutations for each pathway.

### SEM analysis

To quantitatively investigate the interactions among multiple pathways, we conducted a SEM analysis. Because factor analysis in SEM does not allow for identical variables, we temporarily excluded pathways with identical ssGSVA scores from the analysis. First, we used SPSS software (V26.0) to determine the relationship between pathways and factors. Specifically, we obtained the factor loading matrix using principal component analysis (PCA). The number of factors extracted was based on those with eigenvalues greater than 1. We examined the factor loading coefficients in the rotated factor matrix, and variables with inadequate loadings were removed. Specifically, variables were deleted if their loadings on different factors were all below 0.6. This deletion process was repeated until all variables had meaningful associations with the factors. Additionally, factors with only one variable were removed.

Subsequently, the model was constructed and evaluated using LISREL software (V8.7). The model was built based on the Spearman correlation coefficient matrix between pathways and the correspondence between pathways and factors. Modification indices were used to guide the model development. Model improvements were made by considering these indices and the significance of loadings. Specifically, the standard error of the loading of the pathway within their respective factors was considered significant (generally t value >2), and the modification indices for the pathways in other factors remained relatively low.

### Validation with literature evidence

We manually retrieved and summarized AD-related pathways from the literature that were validated by high-throughput data analysis or experiments. These correlations were classified as direct, indirect, or unrelated. “Direct” refers to a direct association of the pathway or factors within the pathway with AD. “Indirect” means the pathway or its factors are correlated with a specific feature that is linked to AD. Other pathways were deemed unrelated to AD.

## Acknowledgements

Authors would like to thank the Dr. Nilüfer Ertekin-Taner who leads MayoRNAseq study. The results published here are in whole or in part based on MayoRNAseq data obtained from the AD Knowledge Portal. We also thank the Supercomputer Center of Chongqing Medical University for their computing power and technical support. This research is financially supported by the Science and Technology Research Program of National Natural Science Foundation of China (Grant No. 32200446), CQMU Program for Youth Innovation in Future Medicine (Grant No. W0147), and Chongqing Municipal Education Commission (Grant No. KJQN202100402).

## Data availability

The data used in this study include MayoRNAseq whole-genome sequencing variant call formats (https://www.synapse.org/#!Synapse:syn11724002), models of gene expression for use in GReX (https://zenodo.org /records/3842289#.YrvrM7FBVYA), and genome-wide summary statistics: (https://ctg.cncr.nl/software/summary_statistics/).

We used publicly available software for all analyses, including Plink 1.9 (https://www.cog-genomics.org/plink2/), PRScs (https://github.com/getian107/PRScs), and MetaXcan (https://github.com/hakyimlab/PrediXcan and https://github.com/hakyimlab/MetaXcan/wiki/Individual-level-PrediXcan:-introduction,-tutorials-and-manual)

Our framework are stored in Github: https://github.com/BFGBgroup/Pathway/tree/main.

